# MetaFunc: Taxonomic and Functional Analyses of High Throughput Sequencing for Microbiomes

**DOI:** 10.1101/2020.09.02.271098

**Authors:** Arielle Kae Sulit, Tyler Kolisnik, Frank A Frizelle, Rachel Purcell, Sebastian Schmeier

**Affiliations:** School of Natural Sciences, Massey University, Auckland, New Zealand; Department of Surgery, University of Otago, Christchurch, New Zealand; Evotec SE, Hamburg, Germany

**Keywords:** metatranscriptomics, microbiome, functional annotation, host correlation

## Abstract

The identification of functional processes taking place in microbiome communities augment traditional microbiome taxonomic studies, giving a more complete picture of interactions taking place within the community. While there are applications that perform functional annotation on metagenomes or metatranscriptomes, very few of these are able to link taxonomic identity to function or are limited by their input types or databases used. Here we present MetaFunc, a workflow which takes input reads, and from these 1) identifies species present in the microbiome sample and 2) provides gene ontology (GO) annotations associated with the species identified. In addition, MetaFunc allows for host gene analysis, mapping the reads to a host genome, and separating these reads, prior to microbiome analyses. Differential abundance analysis for microbe taxonomies, and differential gene expression analysis and gene set enrichment analysis may then be carried out through the pipeline. A final correlation analysis between microbial species and host genes can also be performed. Finally, MetaFunc builds an R shiny application that allows users to view and interact with the microbiome results. In this paper, we showed how MetaFunc can be applied to metatranscriptomic datasets of colorectal cancer.

## Background

Metagenomic or metatranscriptomic studies of microbiome communities allow for characterization of functional contributions as well as taxonomic load, by allowing the identification and quantification of genes possibly contributed by the microbial community. The ability to identify functional processes from the microbiome gives a more complete picture of microbe-microbe and/or microbe-host interactions that drive community dynamics (1).

There are existing bioinformatics programs (2–4) that perform functional annotation on metagenomes and metatranscriptomes, but most of these are unable to link taxonomies (the microbes under study) to their respective functional processes. Existing packages with this capacity include PICRUSt and PICRUSt2 (5,6), and HUMAnN2 (7). PICRUSt and PICRUSt2 predict metagenome function by inferring genes present in OTUs based on their phylogenetic similarities to other OTUs with known gene content (5,6). However, they do not directly measure the genes involved, but rather rely on 16S gene marker sequences, which, being highly conserved, are useful for the identification of bacterial genera (8,9) and are not present in other microbes aside from Bacteria and Archaea (10). Thus 16S based taxonomic identification, and subsequent functional predictions, may be unsuitable for species level identification, and for recognizing other microbes aside from Bacteria and Archaea. HUMAnN2’s taxonomic profiling, meanwhile, is reliant on MetaPhlAn2 (11,12), which uses clade-specific marker genes from reference genomes. Benchmarking efforts by Ye et al., 2019 (10) highlights the limitations of using the MetaPhlAn2 package that includes non-customizable databases and marker gene-based databases, which results in relatively lower precision and recall in its classification.

To augment such meta-omic studies, we present here a simple, straight-forward pipeline named MetaFunc, a snakemake workflow (13) that maps function to a microbiome (and optionally host) sample. MetaFunc uses Kaiju (14) as its main taxonomic classifier. Kaiju uses protein translations of input reads to generate taxonomic profiles. By generating protein-based classifications using metatranscriptomic reads, MetaFunc is able to determine which microbes are more metabolically active, allowing more focus on the functional contributions of microbes. MetaFunc then uses protein accession numbers from Kaiju results to obtain the set of Gene Ontology (GO) terms associated with the microbiome community. Furthermore, Kaiju outputs provide a direct protein – taxonomy ID relationship that makes it possible for MetaFunc to establish which organisms are contributing to the functional GO terms. MetaFunc also has options for pre-processing of reads before running Kaiju: trimming of input reads with fastp (15) can be performed in addition to pre-mapping to a host genome (e.g. human) using STAR (16). The unmapped reads following STAR processing are the input used by MetaFunc for microbe identification, while host gene expression information can be obtained from STAR-mapped reads. Thus, MetaFunc allows simultaneous investigation of host and microbe community active functional processes, as well as active host genes and microbes.

### Protocol

#### Workflow

**Figure 1** shows the workflow that takes place within MetaFunc. Paired-end and/or single-end sequencing reads are used as input in fasta or fastq format. If trimming and mapping are not enabled, reads are used as input to Kaiju and subsequent microbiome analyses (**Figure 1a**). If trimming is enabled, reads are trimmed for adapters and quality controls using fastp. If mapping is enabled, either the trimmed reads or raw input reads are first mapped to a designated host genome using STAR. Unmapped reads after host mapping are then used as input to Kaiju. STAR results are then used to obtain host gene information (**Figure 1b**).

**Figure 1.**
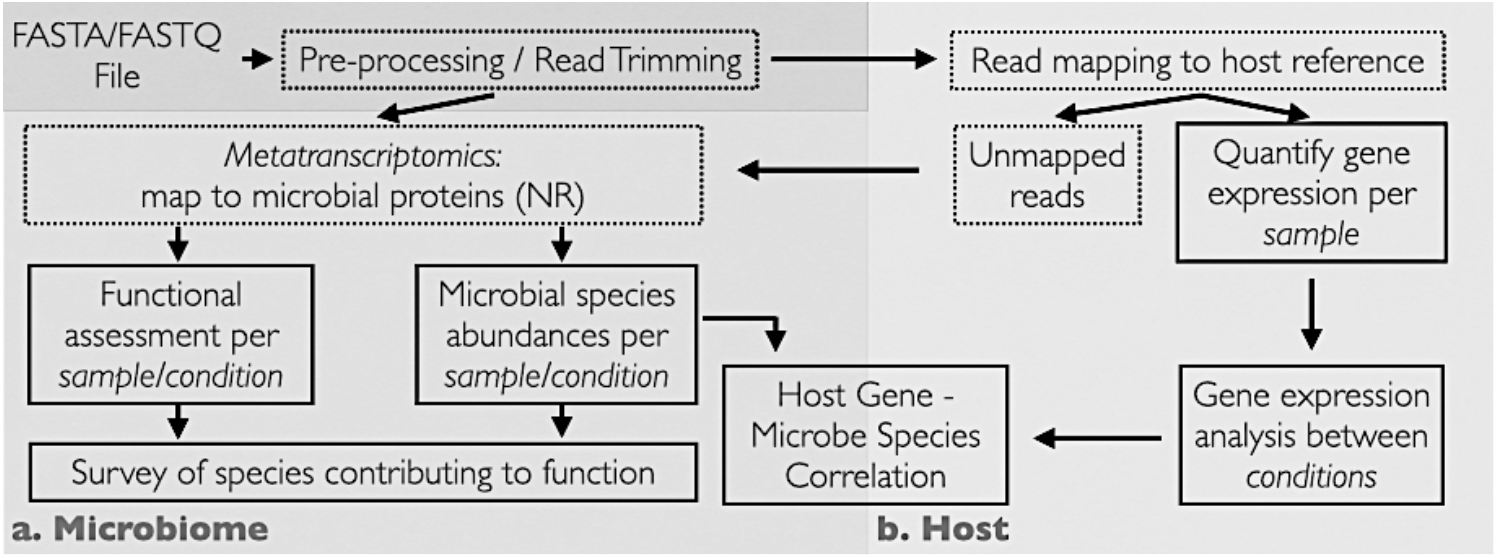
MetaFunc Workflow. The workflow uses FASTQ or FASTA as input and processes reads through the microbiome pipeline to give microbial abundance and function (**a**) and/or host gene analysis (**b**) which will first map reads to a host before sending unmapped reads to the microbiome pipeline. Applying host read analysis will give gene expression analysis results as well as host gene-microbial species correlation. Solid boxes indicate steps with an output while dotted boxes indicate intermediate steps in the pipeline. ***NR: NCBI Blast nr database***

#### Microbiome Analysis (Figure 1a)

MetaFunc parses through Kaiju results and gathers taxonomy IDs of species for taxonomic characterization per sample and their corresponding protein accession numbers, which are subsequently annotated with gene ontology terms.

##### Taxonomy

Each read matches to a taxonomy ID in Kaiju. MetaFunc gathers the species level matches and adds up the raw reads matching to each species taxonomy ID. In cases of strain level identification, MetaFunc adds this count to its parent species. It also obtains scaled read counts in percentages by dividing the final read count of each taxonomy ID by the total reads that have mapped to species level taxonomies. For a dataset, the pipeline removes any taxonomy ID that is less than 0.001% in abundance in all samples of the dataset; this filter removes thousands of species that are likely to be false positives while retaining more confident classifications. Any remaining false classifications are thought not to affect downstream analyses, as the levels would be too low to impact true abundance (10), however, this value can be adjusted by the user. The taxonomy IDs that have passed the cutoff are then used in subsequent analyses. It should be noted that the pipeline still uses the original scaled percent abundances even after filtering. The pipeline would also include the lineage of the taxonomies using *TaxonKit* (17).

For a dataset, the MetaFunc pipeline outputs two tables containing species as rows and samples as columns with values being raw read counts or percent abundance for each species in the samples. If the user wishes to compare groups or conditions (e.g. disease state vs control), the pipeline calculates the average percent abundance of species among samples belonging to a group and this table is also given as an output. Differential abundance of microbes between groups is also carried out in MetaFunc using edgeR (18,19). Raw read count tables are first filtered using the function *filterbyExpr* with threshold of 1 which is user-adjustable, and normalization factors are calculated by *calcNormFactors* with default settings. *exactTest* is then applied to calculate differential abundance with *p*-values adjusted using Benjamini & Hochberg correction or False Discovery Rate (FDR).

##### Proteins

Kaiju outputs the accession number(s) of the protein match(es) with the highest BLOSUM62 alignment score of the read after translation into 6 open reading frames (ORF). It is possible to have more than one best protein match if two or more protein matches have equal scores in Kaiju. In order to account for this, we use proportional read counts per protein accession number where one read is divided by the number of best protein matches it has. Similar to that for taxonomy IDs, the pipeline adds up the proportional read counts per protein accession number of a species. Scaled reads as percent abundances are obtained by dividing the proportional count of each accession number by the total read counts that have mapped to a species.

##### Gene Ontology: Database Construction

Metafunc relies on Kaiju’s nr_euk database for its taxonomic identification and corresponding protein matches. The nr_euk database is built on a subset from NCBI BLAST *nr* database containing Archaea, Bacteria, Fungi, Viruses, and other Microbial Eukaryotes (see https://raw.githubusercontent.com/bioinformatics-centre/kaiju/master/util/kaiju-taxonlistEuk.tsv). Identical sequences in the *nr* database are compiled into 1 entry and Kaiju only outputs the first protein accession number of an entry that has multiple identical sequences (14). Thus we needed to construct the protein-to-GO database such that all functional terms of any protein compiled in 1 *nr* entry are considered.

To facilitate gene ontology annotations, we constructed an sqlite database in which GO annotations of a protein accession number from Kaiju can be looked up. We first gathered relevant NCBI *nr* database entries, converted all of the proteins of an *nr* entry into UniProt (20,21) entries, and then gathered corresponding GO annotations using the Gene Ontology Annotation (GOA) database for all those proteins (22). All GO annotations of one *nr* entry are then linked to the first protein of that entry in an sqlite database, which is used to annotate Kaiju protein accession matches with GO IDs. For more detailed information, please see the Notes section of the pipeline’s documentation page (https://metafunc.readthedocs.io/en/latest/notes.html). For MetaFunc, we provide pre-made databases for download (23,24) but users can make their own updated databases following instructions from https://gitlab.com/schmeierlab/metafunc/metafunc-nrgo.git.

##### Gene Ontology: Protein Annotation

For each sample, the pipeline obtains only the proteins that are from taxonomy IDs that passed cutoffs in the *Taxonomy* section described above. Their scaled proportional read counts, as in the *Proteins* section above, are still scaled against the total number of reads that mapped to a species. In order to compare groups or conditions, the pipeline first calculates the average of the corresponding proportional reads and scaled proportional reads of a protein accession number among samples of a group. It then searches for the GO terms annotating the (*nr*) protein using the created sqlite database described in *Gene Ontology: Database Construction*. Each GO term set annotating an accession number is then updated by accessing parent terms related to the GO terms by *‘is_a’* or *‘part_of’* using *GOATOOLS* (25). Note that this update takes the entire set of GOs annotating the accession number into consideration such that no GO terms or path/s to the top of the GO directed acyclic graph (DAG) is doubled. GOATOOLS also parses other information regarding the go term such as description, namespace, and depth through the go-basic.obo file (26). The proportional and scaled read counts are then added to all GO terms annotating a protein, including updated terms. Finally, the percentage of reads covering a GO term within a namespace (Biological Process, Molecular Function, and Cellular Component) is calculated by dividing the scaled read count of a GO term by the total scaled read counts covering a namespace and multiplying by 100. The final output table of the pipeline is a contingency table with GO IDs of all namespaces as rows and samples or groups as columns, with percentage within a namespace as values.

##### Visualization

To facilitate exploration of results from MetaFunc, MetaFunc automatically builds an R shiny application, such that users can view and interact with the taxonomy and gene ontology tables. The application allows users to select GO terms and identify the species whose proteins are annotated with the searched for term. Conversely, users may search for a species and obtain all GO terms associated with the searched for species. See the pipeline’s documentation page for more information (https://metafunc.readthedocs.io/en/latest/rshiny.html).

#### Host Analyses (Figure 1b)

Many microbiome communities are often associated with a host genome. Reads belonging to the host genome have the capacity to misclassify as microbiome (10) and filtering of host reads has been a part of many microbiome studies, either prior to sequencing or *in silico* (27–29). The MetaFunc pipeline offers the option of mapping reads to a host genome using the program STAR and using the unmapped reads from this step as input to Kaiju for the microbiome analysis.

MetaFunc also allows additional analyses of host reads after STAR mapping. Host genes are quantified using featureCounts (30) of the subread package. If comparisons between groups are indicated, edgeR is used to perform differential gene expression analysis (DGEA). Additionally, supplying a gene matrix transposed (.*gmt*) file from e.g. the molecular signatures database (31–33) allows for gene set enrichment analysis (GSEA) of host genes using the clusterProfiler package (34).

#### Host Gene – Microbe Species Correlation

When a comparison between groups is specified, the pipeline also performs Spearman correlation analysis between the top most significant differentially expressed genes (DEGs) and top most significant differentially abundant (DA) microbes. Results of these correlations are summarized in a matrix on which hierarchical clustering is performed and a heatmap is generated using Clustergrammer (35). Through this heatmap and table, a user can investigate the strength of correlation (*rho*) between a DA microbe and a DEG, and which microbes and genes have similar patterns of correlations.

#### Tutorial/Manual

For a more detailed description of the workflow, usage instructions, and results, documentation of the MetaFunc pipeline may be found in https://metafunc.readthedocs.io/en/latest/index.html.

### Validation

#### Dataset PRJNA413956: Matched Colorectal Cancer (CRC) and Adjacent Non-Tumor Tissue

In order to demonstrate the utility of the MetaFunc pipeline, we obtained publicly available transcriptomics data from the study of Li et al., (2018) (36) consisting of 10 tumor and corresponding adjacent non-tumor colorectal tissue samples. Raw sequencing data was downloaded from https://www.ncbi.nlm.nih.gov/geo/query/acc.cgi?acc=GSE104836 and input to the pipeline and the full workflow carried out, generating data for host, microbiome, and host-microbiome correlation.

##### Microbiome Results

###### Taxonomy

The MetaFunc pipeline outputs a table of percent abundances of species that are identified in each sample and an average of these abundances across members of the same group if a grouping condition is applied. We ran the pipeline with the intent of comparing microbiome species and function between colon cancer samples and non-tumor matched samples.

Previous studies have already established that certain microbes associate more with colorectal cancer samples compared to healthy controls. We searched for *Fusobacterium nucleatum, Parvimonas micra*, and *Porphyromonas asaccharolytica* in the averaged group results. These microbes have previously been found to be more abundant in colorectal cancer cohorts in meta-analyses of several datasets (37,38). We also searched for *Bifidobacterium* species, *B. bifidum* and *B. longum*; *Bifidobacteria* are thought to confer protection from colorectal cancer (39).

The bars in **Figure 2A** show the average percent abundance of the species between samples from tumor and matched non-tumor tissue as identified through MetaFunc. As MetaFunc provides a per sample data, we are also able to plot individual values of CRC (red) and matched normal (blue) samples.

**Figure 2.**
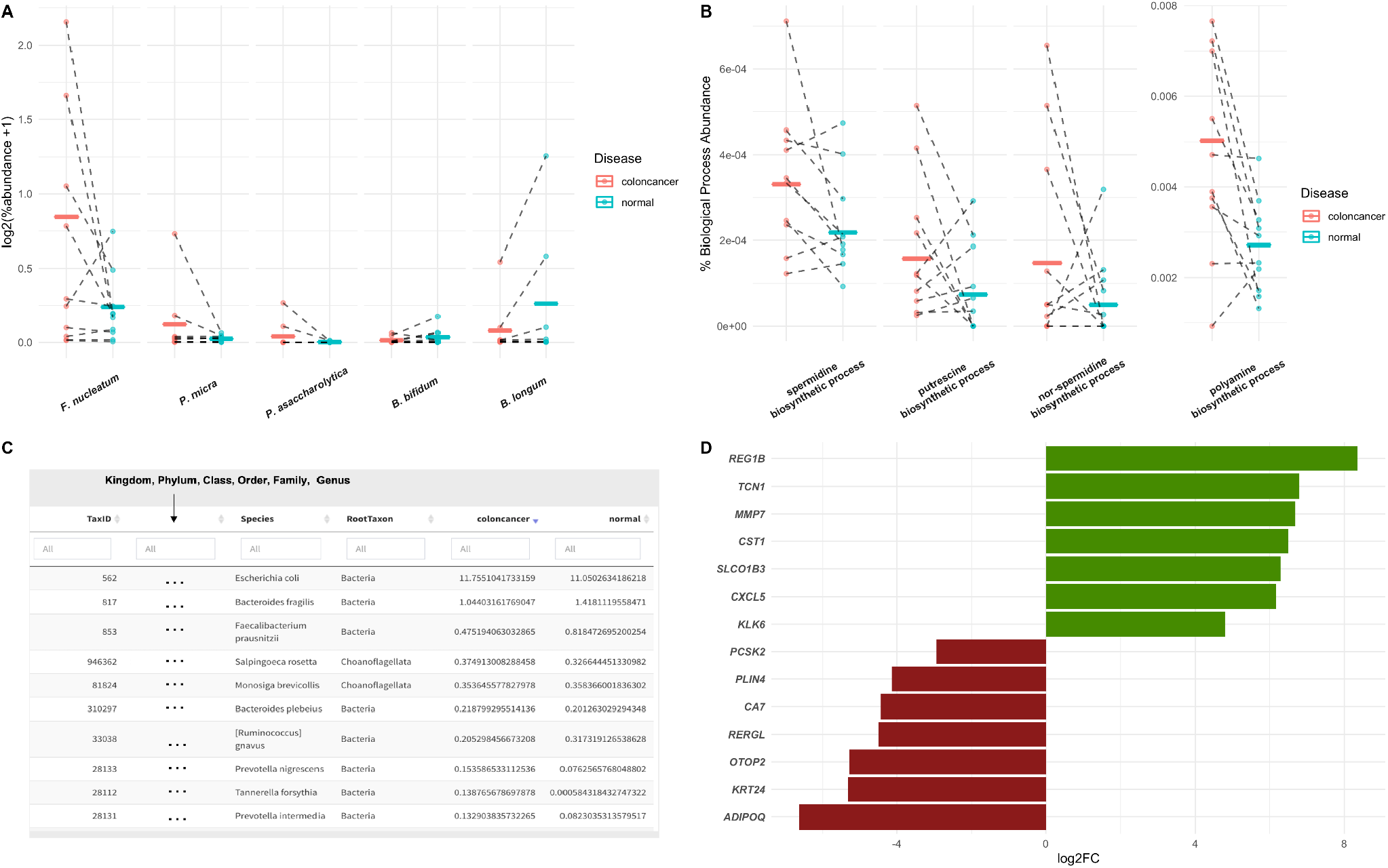
MetaFunc Microbiome and Host Analyses of Dataset PRJNA413956. **A. Average percent abundance of selected bacterial species in CRC tissue compared to matched non-tumor (normal) samples.** From MetaFunc tabulated results, we plotted the percent abundances of selected bacteria in CRC and matched normal samples. Raw values were first log_2_ transformed, with prior addition of 1 as a pseudocount to account for 0 values. Individual points represent per sample transformed values in red (CRC) and blue (Normal). Per group means are represented by the horizontal lines. **B. Percent abundance of specific polyamine biosynthetic process GO terms among all biological process GOs in a sample/group compared between CRC (red) and normal (blue) samples**. Values were calculated as described in section *2*.*3*.*2*.*4 Gene Ontology: Protein Annotation*, and output in MetaFunc tables or in the R Shiny application. These values were plotted, overlaying group means (horizontal lines) and individual values (data points). **Screenshot from MetaFunc R shiny application**. This view shows the first 10 species with proteins contributing to the GO *Polyamine Biosynthetic Process*. The R Shiny application columns include a URL (not shown in screenshot), which is linked to the NCBI’s Taxonomy Browser, the Species Taxonomy ID, Lineage (indicated as ‘*…*’ in screenshot), Root Taxon, and percent abundances of the species in the two groups being compared: CRC and normal samples. Note that percent abundances refer to the total abundance of the species in question, not just the proteins contributing to the GO term. Results can be sorted based on any column from highest to lowest percent abundance in the colon cancer cohort. **D. Fold change of representative upregulated and downregulated human genes** (36) **between CRC and matched normal samples in this study. Fold change values were obtained from the edgeR results of the pipeline. All these genes are significant (FDR < 0.05) in both this study and the source publication**.

As seen in **Figure 2A**, MetaFunc identified *F. nucleatum, P. micra*, and *P. asaccharolytica* as being relatively more abundant in the CRC group while the *Bifidobacterium* species are relatively more abundant in the normal group.

MetaFunc also has a step that utilizes edgeR to perform differential abundance on per sample species read counts, stratified according to CRC and non-tumor grouping. This resulted in a total of 117 species that were significantly different between the groups (*FDR < 0*.*05*). There are 59 species upregulated and 58 downregulated in colon cancer samples. Through the MetaFunc results, we identified *Tanerella forsythia* as the most prominent enriched species in the colon cancer cohort with a *log*_*2*_ *FC = 7*.*40. T. forsythia* is a known oral pathogen, thought to be part of the so-called Red complex of periodontal pathogens, along with *Porphyromonas gingivalis*, and Treponema denticola (40). Members of this Red Complex have been found to be enriched in subtype CMS1 of colorectal cancers (41), the subtype most associated with immune process activation in CRC (42–44).

###### Function

MetaFunc is intended to enable comparisons of the functional potential of the microbiome between groups. MetaFunc uses gene ontology annotations of protein matches from Kaiju. To demonstrate, we focused on polyamine biosynthetic processes GO terms. Polyamines (PAs) are polycations found to play important biological functions in cell growth. These molecules have been found to be associated with tumor progression and growth (45–47). Although cells are able to biosynthesize polyamines and even export them, a large source of cellular polyamines comes from uptake from their surroundings and, importantly, the microbiota is thought to be an essential source (38,46,47) with spermidine and putrescine being the most common of bacterial PAs (47).

The bars in **Figure 2B** show the percent of reads among biological process GOs covering PA biosynthetic processes in the Colorectal Cancer and Normal conditions, superimposed with the individual values of samples from the CRC (red) and Normal (blue) groups. From **Figure 2B**, we saw that several of the polyamine biosynthetic processes were relatively more abundant in the colorectal cancer cohort compared to the normal cohort, using protein annotations.

We used the built-in MetaFunc shiny application to facilitate an inquiry into the microbes species that may contribute to polyamine synthesis. To illustrate, we searched for *‘polyamine biosynthetic process’* in the ‘GO to TaxIDs’ tab of the application, and obtained a total of 126 TaxIDs contributing to the GO term in both colorectal cancer and normal samples. Of these TaxIDs, we identified *E*.*coli* and *B. fragilis* to be most abundant in both cohorts. However, differences in relative abundance of some microbial species can be identified between cancer and normal cohorts, notably several of which are oral pathogens from the genus *Prevotella*. A striking difference in abundance was seen in *Tannerella forsythia*, which was previously found to be significantly more abundant in the colorectal cancer cohort via edgeR (**Figure 2C**). These data suggest that *T. forsythia* represents one of the bacterial species that most contributes to increased polyamine synthesis in CRC samples in this cohort.

###### Host Results

The dataset we used for this study was from a total RNA transcriptomics run aiming to identify long non-coding RNAs (lncRNAs) and mRNAs in colorectal cancer samples (36). Therefore, we first mapped the reads to the human genome using the STAR mapping utility of the pipeline, subsequently using only the unmapped reads for the microbiome analyses. From the reads mapped to the human genome, MetaFunc was able to obtain counts of reads covering human genes and using these, obtained differentially expressed genes between CRC and matched normal samples through edgeR. MetaFunc results showed a total of 1476 differentially expressed genes with an *FDR < 0*.*05* and |log_2_fold change|>2. From these, we found all the top 5 upregulated and top 5 downregulated genes as reported in the source publication (36), as well as all the genes they had randomly selected for expression confirmation via qPCR. **Figure 2D** shows their fold change as found through MetaFunc.

MetaFunc is also able to perform host gene set enrichment analysis using the differentially expressed genes. Significant gene sets (*p*.*adjust < 0*.*05*) with the highest normalized positive enrichment scores (NES) included such terms as ribosome biogenesis, DNA replication, mitotic nuclear division, and condensed chromosome (see **Supplementary Table 1**), many of which appear to be related to cell division or replication, consistent with the findings of the source publication (36), that the upregulated lncRNAs they found were involved in mitosis, cell cycle process, and mitotic cell cycle.

###### Host – Microbiome Correlations

We set MetaFunc’s default abundance cutoff for microbial identification to 0.001% to remove most probable contaminants and so as not to lose any other meaningful taxonomies. It has been shown in a prior study (10), however, that most classifiers call false positives at below 0.01% abundance. We therefore applied this 0.01% cutoff in looking at the host-microbiome correlations in this dataset to narrow our focus on microbes that are more likely to be involved in our test case.

In using the 0.01% cutoff, MetaFunc was able to only identify 19 differentially abundant microbes. Their correlations with the top 100 significantly abundant genes can be seen at the URL: http://amp.pharm.mssm.edu/clustergrammer/viz/5f02a49e8ec9bb33170b865c/cor.deg-tax.matrix.tsv. **Table 1** highlights some notable correlations between differentially abundant microbes and differentially expressed human genes. *T. forsythia*, although significantly abundant in CRC samples, do not correlate significantly with any DEGs in CRC. Among its highest correlations however included the gene Colorectal Neoplasia Differentially Expressed (CRNDE).

**Table 1.** Spearman Correlation Between DA Microbes and DGEs in CRC.

| Table 1. Spearman Correlation Between DA Microbes and DGEs in CRC |  |  |  |  |  |
| --- | --- | --- | --- | --- | --- |
| Gene Name | Gene ID | TaxID | Species | rho | p-value |
| CRNDE | ENSG00000245694.10 | 28112 | Tannerella forsythia | 0.29 | 0.22 |
|  |  | 36911 | Clavispora lusitaniae | 0.70 | 0.00063 |
|  |  | 106590 | Cupriavidus necator | 0.65 | 0.0019 |
|  |  | 1314 | Streptococcus pyogenes | 0.63 | 0.0027 |
| TCN1 | ENSG00000134827.8 | 106590 | Cupriavidus necator | 0.71 | 0.00042 |
|  |  | 36911 | Clavispora lusitaniae | 0.61 | 0.0045 |
|  |  | 1314 | Streptococcus pyogenes | 0.60 | 0.0048 |
| WNT2 | ENSG00000105989.10 | 1314 | Streptococcus pyogenes | 0.84 | 4.07E-06 |
|  |  | 106590 | Cupriavidus necator | 0.75 | 0.00015 |
|  |  | 36911 | Clavispora lusitaniae | 0.75 | 0.00016 |

Conversely, we investigated which species correlated with CRNDE. The highest correlations were with microbes *C. lusitaniae, C. necator*, and *S. pyogenes*. All correlations were determined to be significant. The same species were among the highest correlations of TCN1, and WNT2. TCN1 was among the top DEGs in cancer identified in this study as well as in the source publication (36). WNT2 meanwhile is part of the *Wnt/β-catenin* pathway, which has roles in cell proliferation, cell migration, and cell differentiation. WNT2 is responsible for hyperactivation of β-catenin and is known to be upregulated in CRC (48).

#### Dataset PRJNA404030: Consensus Molecular Subtypes (CMS) of CRC Samples

To illustrate MetaFunc’s capacity to compare more than two sample groups, we used MetaFunc to analyze transcriptome reads from the study of Purcell and colleagues (41) (raw reads may be accessed at https://www.ncbi.nlm.nih.gov/bioproject/PRJNA404030), which are grouped into four CRC consensus molecular subtypes (CMS). A total of 33 samples were collected during surgical resection of tumors, and sample preparation for RNA sequencing was carried out using the Illumina TruSeq Stranded Total RNA Library preparation kit. For this sample SolexaQA++ (49) was used to trim reads, which were then run through Salmon (50) to quantify transcript expression. The publicly available CRC CMS classifier (43) was used to categorize samples into one of four CMSs. Of the 33 samples, only 27 were classified into a CMS and of these, only one sample was classified into CMS4. This sample was also removed from the dataset for lack of replicates leaving a total of 26 samples – 7 samples in CMS1, 11 in CMS2, and 8 in CMS3. Metafunc was used with default parameters, except for the following options: trimming was set to false, and featureCounts with reverse stranded option was used.

##### Microbiome Results

###### Taxonomy

MetaFunc performed pairwise differential abundance analysis on the three groups using edgeR. From MetaFunc’s results, we considered a species to be significantly abundant in a subtype if it is significantly abundant compared to both of the other subtypes. For instance, a significantly abundant species in CMS1 must be significantly abundant in the CMS1 vs CMS2 and CMS1 vs CMS3 comparisons. Using this definition, only CMS1 had species that were significantly abundant (*FDR <0*.*05*) compared to both CMS2 and CMS3. **Figure 3A** shows the False Discovery Rate (FDR; diamonds) and log_2_ fold change (bars) of the species in CMS1 compared to CMS2 (blue) and CMS3 (brown).

**Figure 3.**
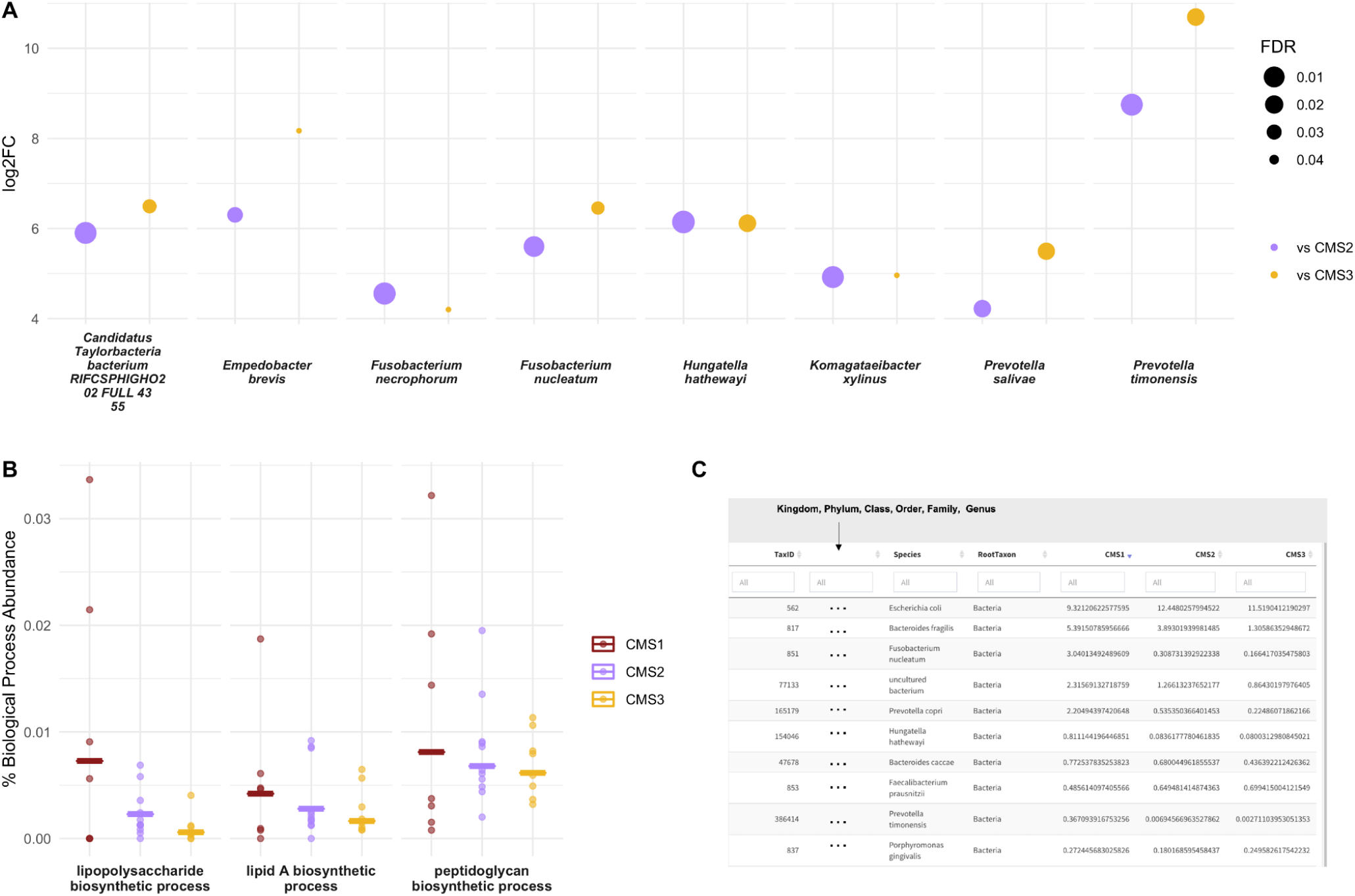
MetaFunc Microbiome Analysis of Dataset PRNJA4040030. **A. Microbes that are significantly more abundant (FDR < 0.05) in CMS1 compared to CMS2 (purple) and CMS3 (yellow).** Microbes are considered differentially abundant (DA) in CMS1 if it is identified through edgeR as DA in both CMS1 vs CMS2 and CMS1 vs CMS3 comparisons. Log_2_FC (y-axis) is the log_2_ of the fold-change between CMS1 and the other subtypes (e.g. CMS1/CMS2); FDR (point sizes) is the false discovery rate adjusted p-values. **B. Percent abundance of specific PAMPs biosynthetic process GO terms among all biological process GOs in a sample/group compared between CRC subtypes, CMS1 (red)**, **CMS2 (purple), and CMS3 (yellow)**. Values were calculated as described in section *2*.*3*.*2*.*4 Gene Ontology: Protein Annotation*, and output in MetaFunc tables or in the R Shiny application. These values were plotted, overlaying group means (horizontal lines) and individual values (data points). **C. Screenshot of R shiny application showing the relative abundances of species associated with PAMPs biosynthetic processes compared among CMS1, CMS2, and CMS3**. This view shows the first 10 species, with highest abundances in CMS1, with proteins contributing to any of the PAMPs biosynthetic processes described above. The application columns show a URL (not shown in screenshot), which is linked to the NCBI’s Taxonomy Browser, the Species Taxonomy ID, Lineage (shown as ‘…’ in screenshot), Root Taxon, and percent abundances of the species in the three groups being compared: CMS1, CMS2, and CMS3. Note that percent abundances refer to the total abundance of the species in question, not just the proteins contributing to the GO term. Results shown are sorted from highest to lowest percent abundance in the CMS1 group.

We take note of species in the genera *Prevotella* and *Fusobacterium*, which have previously been associated with colorectal cancer. *Fusobacterium nucleatum* in particular has strong evidence of an association with CRC (37,51,52). Most of these are also members of the oral microbiota, which have also previously been associated with cancer development particularly through inflammatory processes (53). We found no species that were significantly abundant in CMS2 or CMS3 using the given criteria.

###### Function

Through the microbiome functional results of MetaFunc, we then investigated if processes relating to pathogen-associated molecular patterns (PAMPs) were contributed by the microbial communities, considering that CMS1 is characterized by immune responses, which are usually triggered when the human immune system recognizes such molecules. We used the MetaFunc R shiny application to search for terms *‘lipopolysaccharide biosynthetic process’, ‘lipid A biosynthetic process’*, and *‘peptidoglycan biosynthetic process’*, and their relative abundances. Unsurprisingly, all PAMPs were relatively more abundant in CMS1 (**Figure 3B**).

Using the MetaFunc R shiny application, we also searched for which species might be contributing to the above terms. **Figure 3C** is a screencap of the application showing the species contributing to any of the terms in **Figure 3B. Figure 3C** is arranged from highest to lowest relative abundance in CMS1 and we saw microbes that were among those identified to be significantly abundant in CMS1 such as *Fusobacterium nucleatum, Prevotella timonensis*, and *Hungatella hathewayi*.

###### Gene Set Expression Analysis

MetaFunc calculated differentially expressed genes between subtypes in a pairwise manner (i.e. CMS1 vs CMS2, CMS1 vs CMS3, CMS2 vs CMS3). From the DEGs of the results, MetaFunc was also able to calculate enriched gene sets for each comparison. Similar to identifying DA microbes, we obtained a final set of enriched gene sets for a subtype if it showed enrichment compared to both other subtypes (*p*.*adjust < 0*.*05*). Unsurprisingly, we saw several GO terms involved in immune response enriched in CMS1, including regulation of innate immune response, response to interferon gamma, and positive regulation of cytokine production among others. Enriched GOs in CMS2 are involved in the cell cycle and and ribosome biogenesis, with terms such as tRNA metabolic process, ribosomal large subunit biogenesis, and DNA replication initiation, while GOs enriched in CMS3 involve metabolic processes, e.g. primary xenobiotic metabolic process, flavonoid metabolic process, and lipid catabolic process. These results are consistent with the description of these three CRC subtypes in the original CMS study (43). The top enriched gene sets for each subtype can be found in **Supplementary Tables 2-7**.

##### Host – Microbiome Results

Next, using correlation results from MetaFunc, we investigated which of the top significantly differentially expressed genes correlated with the significantly abundant microbes in CMS1. We obtained the following statistically significant correlations between host and microbiome abundances shown in **Table 2**.

**Table 2.** Spearman Correlation Between DA Microbes in CMS1 and DGEs in CMS1

| Gene Name | Gene ID | TaxID | Species | rho | p-value |
| --- | --- | --- | --- | --- | --- |
| WARS1 | ENSG00000140105.18 | 1802307 | *Candidatus Taylorbacteria bacterium RIFCSPHIGHO2_02_FULL_43_55* | 0.59 | 0.0015 |
| WARS1 | ENSG00000140105.18 | 851 | *Fusobacterium nucleatum* | 0.55 | 0.0035 |
| RNF213 | ENSG00000173821.19 | 1802307 | *Candidatus Taylorbacteria bacterium RIFCSPHIGHO2_02_FULL_43_55* | 0.54 | 0.0048 |
| ICAM1 | ENSG00000090339.9 | 1802307 | *Candidatus Taylorbacteria bacterium RIFCSPHIGHO2_02_FULL_43_55* | 0.50 | 0.01 |
| RNF213 | ENSG00000173821.19 | 851 | *Fusobacterium nucleatum* | 0.50 | 0.01 |
| PARP14 | ENSG00000173193.15 | 851 | *Fusobacterium nucleatum* | 0.47 | 0.02 |
| PARP14 | ENSG00000173193.15 | 1802307 | *Candidatus Taylorbacteria bacterium RIFCSPHIGHO2_02_FULL_43_55* | 0.47 | 0.02 |
| PARP9 | ENSG00000138496.16 | 851 | *Fusobacterium nucleatum* | 0.46 | 0.02 |
| ICAM1 | ENSG00000090339.9 | 851 | *Fusobacterium nucleatum* | 0.46 | 0.02 |
| SLC15A3 | ENSG00000110446.11 | 1802307 | *Candidatus Taylorbacteria bacterium RIFCSPHIGHO2_02_FULL_43_55* | 0.46 | 0.02 |
| STAT1 | ENSG00000115415.19 | 386414 | *Prevotella timonensis* | 0.44 | 0.02 |
| CD163 | ENSG00000177575.12 | 1802307 | *Candidatus Taylorbacteria bacterium RIFCSPHIGHO2_02_FULL_43_55* | 0.42 | 0.03 |
| PARP14 | ENSG00000173193.15 | 386414 | *Prevotella timonensis* | 0.42 | 0.03 |
| CD163 | ENSG00000177575.12 | 386414 | *Prevotella timonensis* | 0.42 | 0.03 |
| ICAM1 | ENSG00000090339.9 | 386414 | *Prevotella timonensis* | 0.41 | 0.04 |
| PARP9 | ENSG00000138496.16 | 1802307 | *Candidatus Taylorbacteria bacterium RIFCSPHIGHO2_02_FULL_43_55* | 0.41 | 0.04 |
| CD163 | ENSG00000177575.12 | 851 | *Fusobacterium nucleatum* | 0.41 | 0.04 |
| SLC15A3 | ENSG00000110446.11 | 851 | *Fusobacterium nucleatum* | 0.40 | 0.04 |
| STAT1 | ENSG00000115415.19 | 851 | *Fusobacterium nucleatum* | 0.40 | 0.04 |
| PML | ENSG00000140464.20 | 1802307 | *Candidatus Taylorbacteria bacterium RIFCSPHIGHO2_02_FULL_43_55* | 0.39 | 0.05 |
| GBP1 | ENSG00000117228.10 | 28448 | *Komagataeibacter xylinus* | -0.39 | 0.05 |
| CEBPA | ENSG00000245848.3 | 154046 | *Hungatella hathewayi* | -0.41 | 0.04 |
| GNLY | ENSG00000115523.16 | 28448 | *Komagataeibacter xylinus* | -0.43 | 0.03 |

Some of these correlations may be found in http://maayanlab.cloud/clustergrammer/viz/610d8b3c97f268000ea37f41/cor.deg-tax.matrix.tsv. This is the hierarchical cluster obtained when correlating top DA microbes and top DGEs in CMS1 compared to CMS2. It is to be noted that there may be correlations in this clustering that are not found in CMS1 compared to CMS3 and are therefore not reported in **Table 2**.

The Spearman correlations (*rho*) between DA microbes and DGEs were quite small in value (the highest value being ∼ |0.59| between WARS1 and *Candidatus Taylorbacteria bacterium RIFCSPHIGHO2_02_FULL_43_55*). Nevertheless, several of the genes appeared to have relevant function with regards to CRC and immune responses. **Table 3** shows information for genes that correlated with *Fusobacteria* and *Prevotella* species in our analyses. These 2 microorganisms have previously been associated with CRC.

**Table 3.**
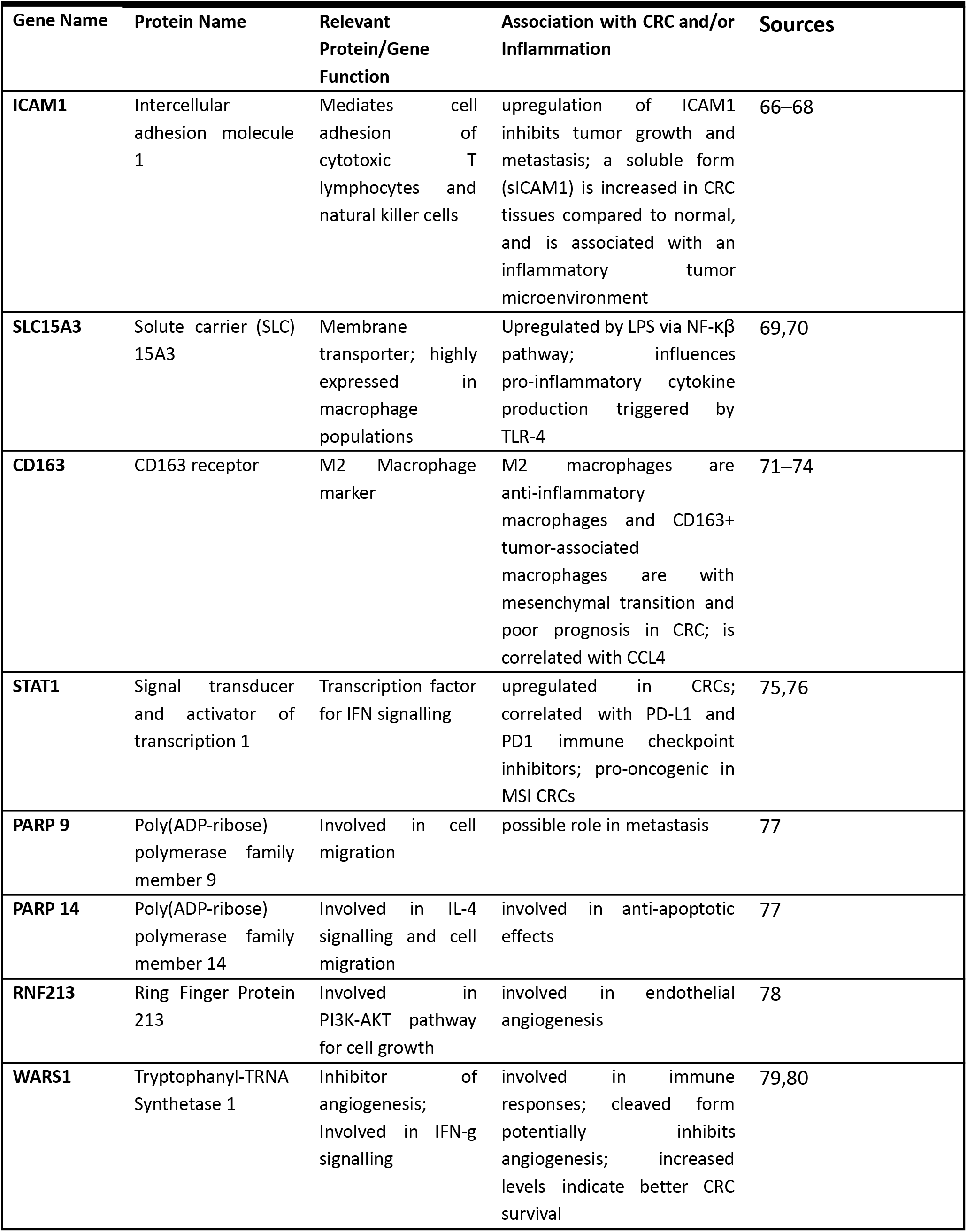
Gene Information of DEGs correlated with DA Microbes in CMS1.

| Gene Name | Protein Name | Relevant Protein/Gene Function | Association with CRC and/or Inflammation | Sources |
| --- | --- | --- | --- | --- |
| **ICAM1** | Intercellular adhesion molecule 1 | Mediates cell adhesion of cytotoxic T lymphocytes and natural killer cells | upregulation of ICAM1 inhibits tumor growth and metastasis; a soluble form (sICAM1) is increased in CRC tissues compared to normal, and is associated with an inflammatory tumor microenvironment | 66–68 |
| **SLC15A3** | Solute carrier (SLC) 15A3 | Membrane transporter; highly expressed in macrophage populations | Upregulated by LPS via NF-κB pathway; influences pro-inflammatory cytokine production triggered by TLR-4 | 69,70 |
| **CD163** | CD163 receptor | M2 Macrophage marker | M2 macrophages are anti-inflammatory macrophages and CD163+ tumor-associated macrophages are with mesenchymal transition and poor prognosis in CRC; is correlated with CCL4 | 71–74 |
| **STAT1** | Signal transducer and activator of transcription 1 | Transcription factor for IFN signalling | upregulated in CRCs; correlated with PD-L1 and PD1 immune checkpoint inhibitors; pro-oncogenic in MSI CRCs | 75,76 |
| **PARP 9** | Poly(ADP-ribose) polymerase family member 9 | Involved in cell migration | possible role in metastasis | 77 |
| **PARP 14** | Poly(ADP-ribose) polymerase family member 14 | Involved in IL-4 signalling and cell migration | involved in anti-apoptotic effects | 77 |
| **RNF213** | Ring Finger Protein 213 | Involved in PI3K-AKT pathway for cell growth | involved in endothelial angiogenesis | 78 |
| **WARS1** | Tryptophanyl-TRNA Synthetase 1 | Inhibitor of angiogenesis; Involved in IFN-g signalling | involved in immune responses; cleaved form potentially inhibits angiogenesis; increased levels indicate better CRC survival | 79,80 |

#### Comparison of Kaiju Results to HUMAnN2

HUMAnN2 (7) is one of the packages most frequently used to assess functional pathways of the microbiome, and to determine which organisms are contributing to the functional pathways. HUMAnN2 works by pre-screening which taxonomies are present in a sample using MetaPhlAn2, afterwards aligning the reads to pangenomes of the classified taxonomies for gene hits. Unclassified reads then undergo an organism-agnostic translated search (7). MetaPhlAn2 has a rather limited database for pre-screening of organisms (10), resulting in a high level of unmapped reads and a limited number of organisms identified.

We ran the same sequencing reads from the study PRJNA413956 (36) through HUMAnN2, first trimming with fastp and removing human-mapped reads using the same conditions as for the MetaFunc pipeline. To be more comparable, we changed the pre-screen threshold of HUMAnN2 to 0.001% of mapped reads. Part of HUMAnN2’s tiered search uses Diamond (7), which requires higher memory and run time compared to Kaiju, used by MetaFunc (10). From taxonomy identification, using Kaiju, to the generation of gene ontology tables, took MetaFunc 11.39 hours to complete, while a comparable analysis using HUMAnN2 took 65.9 hours to complete, almost 6x slower than MetaFunc on the same machine (CentOS Linux release 7.9.2009). Runs for HUMAnN2 may be accessed at https://github.com/asulit08/Humann2_PRJNA413956.

Results showed that for the 20 samples analyzed, 8.4% – 22.9% of reads mapped after nucleotide and protein alignment steps. In contrast, using Kaiju in the MetaFunc pipeline resulted in 33.8% – 56.2% reads mapped to microbial species through protein matches (**Figure 4**).

**Figure 4.**
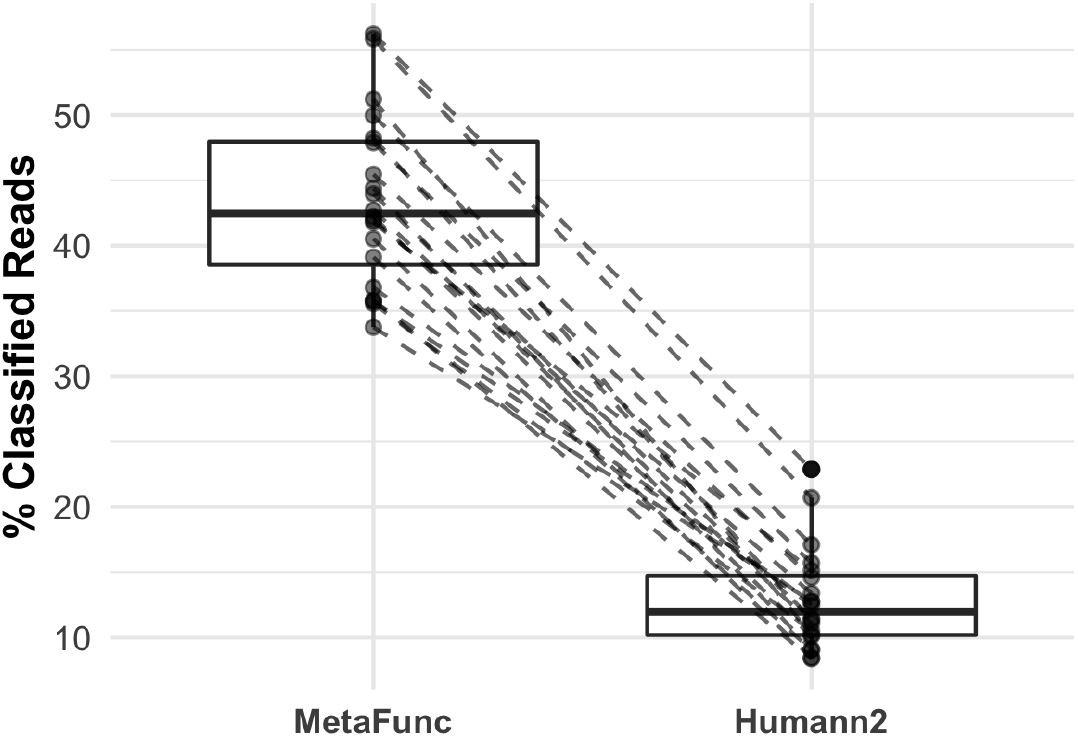
Percentage (%) of reads classified using HUMAnN2 and MetaFunc pipeline using Kaiju as distributed across 20 samples. This data shows that many more reads are classified to a microbiome taxonomy using MetaFunc’s Kaiju compared to HUMAnN2’s MetaPhlan2. Dashed lines connect the same samples as run by either MetaFunc or HUMAnN2.

We also detected only 87 species across the 20 samples using HUMAnN2, compared with a total of 4267 species using Kaiju in the MetaFunc pipeline.

We investigated the same species and polyamine (PA) biosynthetic process GO terms in our HUMAnN2 results as we had in the MetaFunc run of dataset PRJNA413956 (**Supplementary Figures 1 and 2**). We see in **Supplementary Figure 1** that abundances of the species in CRC and normal groups have the same trends in HUMAnN2 results as in that of MetaFunc (Kaiju) results. In HUMAnN2 runs, however, we were not able to find *B. bifidum* among the identified species in the PRJNA413956 cohort. Meanwhile, we also see the same trends in the abundances of PA biosynthetic process GO terms in CRC samples compared to matched normal samples in HUMAnN2 runs, as in our MetaFunc run (**Supplementary Figure 2)**, except for nor-spermidine biosynthetic process, which was not seen using HUMAnN2. Differences in abundance values were noted when comparing individual samples, however, direct comparison between HUMAnN2 and MetaFunc is difficult as raw read-counts scaled to species-classified reads are used in MetaFunc, while HUMAnN2 uses reads-per-kilobase (RPK)-based relative abundances.

## Discussion

MetaFunc allowed us to investigate the relative abundances of known CRC – associated bacteria between CRC samples and matched normal tissues using the *PRJNA413956* dataset. MetaFunc results showed that microbes known to contribute to CRC progression are relatively more abundant in cancer samples while those protective in CRC are relatively more abundant in the normal samples. Through MetaFunc, we also identified that *Tannerella forsythia*, a known oral pathogen and part of the Red Complex that causes periodontal diseases (40), is significantly more abundant in CRC tissues than in normal tissues. Oral pathogens have previously been seen to associate with CRC samples (38,53,69,70). By investigating the R shiny application from MetaFunc, we also found that *T. forsythia*, along with bacteria in the *Prevotella* genera, contributed to polyamine biosynthetic processes indicating that some oral pathogens contribute to cancer progression by producing polyamines that could be taken up by the surrounding cells.

Furthermore, we were able to find known bacteria in the MSI-Immune subset of CRCs by identifying the DA microbes in CMS1 compared to both CMS2 and CMS3 subtypes, as identified by MetaFunc’s edgeR step. *Fusobacteria* have long been associated with colorectal cancer development (37,38,51,52) while *Prevotella* includes species that inhabit the oral cavity; there have also been *Prevotella* species that were found to be abundant in CRC cohorts (37,51,69). In line with this, PAMPs were also found to be relatively more abundant in the CMS1 cohort upon investigation through MetaFunc’s R shiny application. The involvement of these bacteria in CMS1 as well as a relatively higher abundance of proteins contributing to biosynthesis of PAMPs in CMS1 indicate a role of microorganisms in the immune responses that drive the development of CRC in these tumors. This is further supported by correlation with host genes involved in inflammation and/or CRC development as found using MetaFunc’s spearman correlation step. The lack of significantly abundant microorganisms in CMS2 and CMS3 may reflect that the CRC development in these subtypes are not as dependent on immune dysregulation.

We created MetaFunc with the aim of identifying microbes and their functional contribution in a microbiome environment. One of the most widely used packages for this is HUMAnN2 (7) but we find the taxonomic identification generated by HUMAnN2 to be limited, because of its reliance on marker genes. Although similar trends were seen in taxa and gene ontologies of interest between CRC and matched normal samples, fewer test reads were designated as taxa using HUMAnN2 compared to MetaFunc in our comparative analysis. Unfortunately, direct comparison was not possible because HUMAnN2 and MetaFunc use different abundance outputs. For our purposes we found MetaFunc necessary for investigating novel microbes that did not have marker gene representation, faster runs for larger amounts of data, and compatibility with downstream analysis programs.

We acknowledge that, especially at the 0.001% abundance cutoff, some of these species we are seeing could be false positives, or that these could be contaminants from sequencing and processing kits used (71,72). We would caution users in interpreting data from microbes of very low abundances and would recommend following the advice of including negative control samples in sequencing (72). Indeed we could be seeing these effects upon looking at the microbes correlating with significantly abundant host genes in CRC samples from *PRJNA413956*. While *C. lusitaniae* is an opportunistic pathogen causing candidemia (73,74) possibly exploiting the lowered immune responses in cancer patients (75), and some *Streptococcus* species have previously been implicated in CRC (76,77), with *S. pyogenes* having been known to cause invasive infections in humans (78), *Cupriavidus necator* (formerly known as *Ralstonia eutropha* (79)), is a soil bacterium that may be a sequencing contaminant in this dataset. *Cupriavidus* and *Ralstonia* species have been previously identified as common contaminants in meta-omics studies (72,80).

MetaFunc analyzes host and microbiome reads, providing a user-friendly, interactive R-shiny application to investigate results, most useful for those with candidate microbes and function in mind, or for exploratory analyses of the characteristics of a user’s dataset. Downstream analysis, such as differential abundance of microbes, can also facilitate parsing of tables in the shiny application. Its results also provide potential starting points for more in-depth analyses or hypothesis generation for experimental procedures. Results in *‘*.*tsv’* formats are also provided for use in other downstream bioinformatics applications.

While this method was developed for a metatranscriptomic dataset, it is also suitable for metagenomic data input. As Kaiju (14) identifies a single best protein match (or multiple matches with equal scores) of a read, we recommend its usage for short-read datasets. An exception could be made for long read sets in which the user is certain an input read will only span one protein.

We used the MetaFunc pipeline to compare genes and microbes between or among groups, but exploratory analyses of datasets from single groups can also be carried out.

## Conclusion

Here we presented MetaFunc, a singular pipeline for analyzing host and microbiome sequencing reads and their relationships. We found that we identified more microbes in our test datasets using MetaFunc compared to HUMAnN2, while microbes and functions of interest were comparable between the two. We have used MetaFunc to determine that microbes previously known to have associations with CRC are indeed relatively more abundant in CRC samples compared to normal samples. Furthermore, we were able to use MetaFunc to highlight that these microorganisms could contribute to CRC progression through polyamine production.

For a dataset with more than two groups, we have also used MetaFunc to identify abundant bacteria in a CRC subtype associated with immune responses, while conversely, we have not been able to identify significant microbes in the other CRC subtypes. MetaFunc’s spearman correlation step showed that the significant bacteria correlate with human DEGs that function in immune responses and CRC progression. We showed that MetaFunc was able to identify candidate microorganisms that differentiate sample groups and provide insight on the functional capacities of these candidates.

## Acknowledgement

The authors would like to thank Dr. Olin Silander for valuable technical and academic advice for this manuscript.

## Code Availability

MetaFunc is freely available through https://gitlab.com/schmeierlab/workflows/metafunc.git, and full documentation can be found in https://metafunc.readthedocs.io/en/latest/.

