## Supplementary Material for "MetaFunc: Taxonomic and Functional Analyses of High Throughput Sequencing for Microbiomes"

### Supplementary Figures

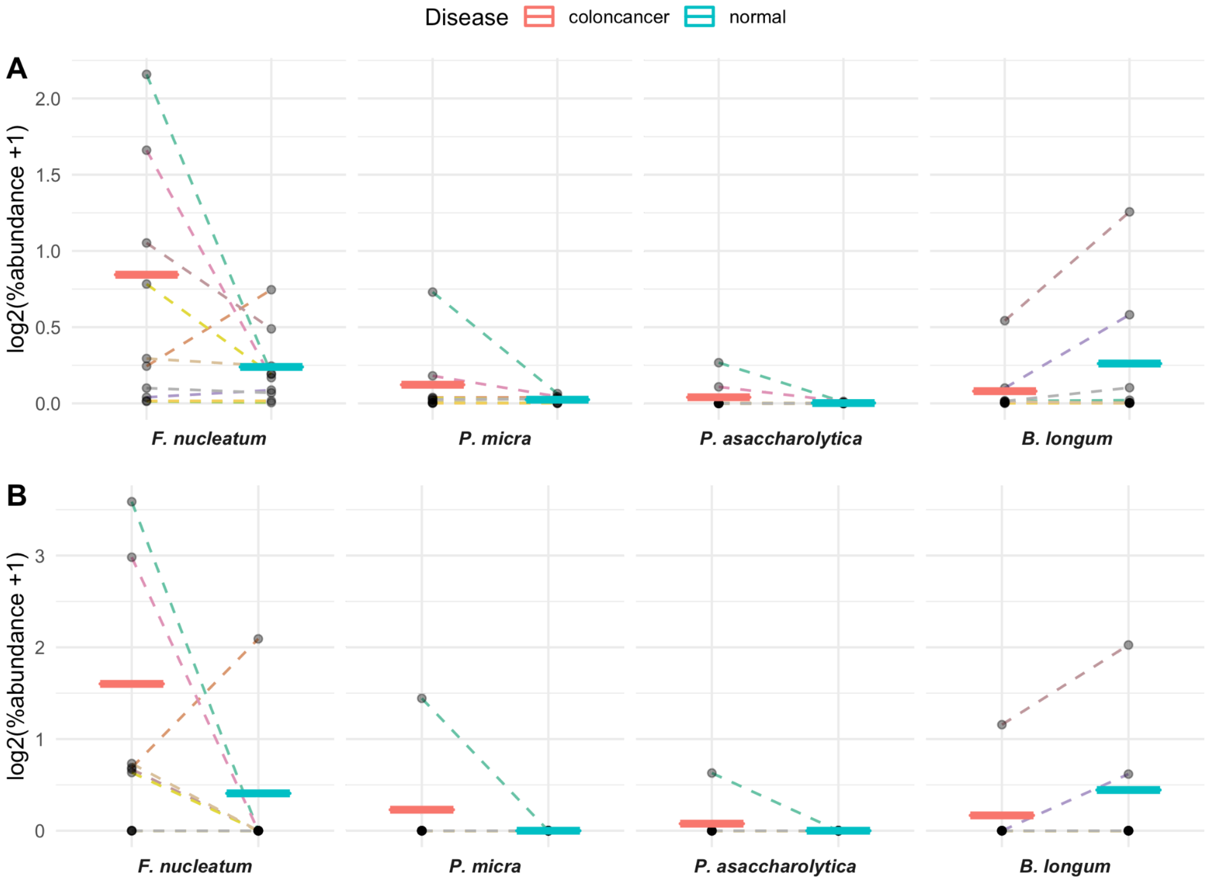


**Supplementary Figure 1. Average percent abundance of selected bacterial species in CRC tissue compared to matched non-tumor (normal) samples as measured by A.** MetaFunc (top) **or B.** HUMAnN2 (bottom)**.** We plotted the percent abundances of selected bacteria in CRC and matched normal samples. Raw values were first $\log_2$ transformed, with prior addition of 1 as a pseudocount to account for 0 values. Individual points represent individually transformed sample values. Per group means are represented as horizontal lines and colored according to disease state. Dashed lines connect matched CRC and normal values. Line colors correspond to individual patients.

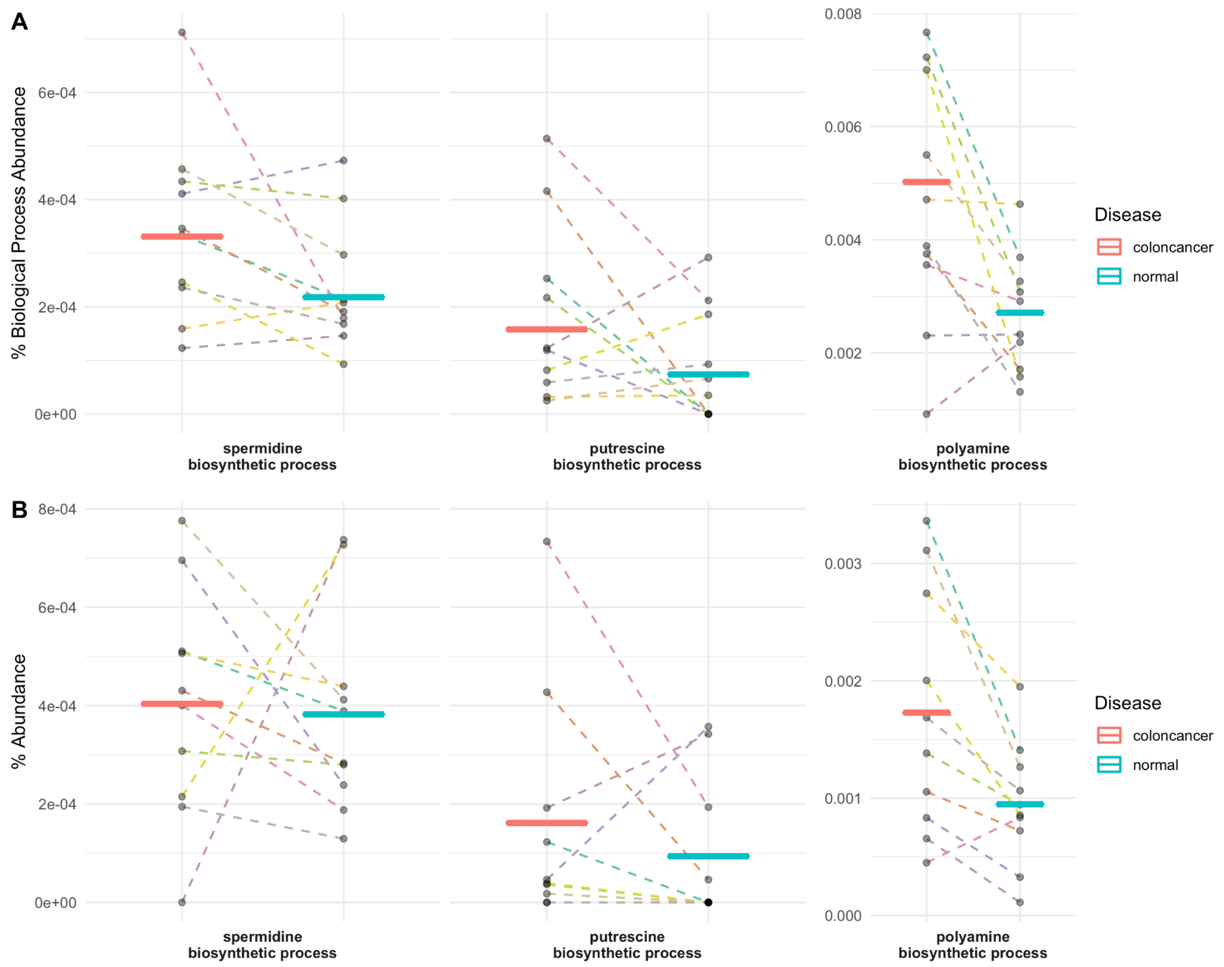


**Supplementary Figure 2. Polyamine biosynthetic process GO terms compared between CRC (red) and normal (blue) samples. A. As measured by MetaFunc.** Percent abundance of specific polyamine biosynthetic process GO terms among all biological process GOs in a sample/group. Values were calculated as described in section *2.3.2.4 Gene Ontology: Protein Annotation,* and output in MetaFunc tables or in the R Shiny application. These values were plotted, overlaying group means (horizontal lines) and individual values (data points). Line colors depict samples from a single patient. Lines connect paired CRC and normal samples. **B. As measured by HUMAnN2.** Percent abundance of specific polyamine biosynthetic process GO terms. HUMAnN2 outputted read counts as reads-per-kilobase (RPKs), which was then converted to counts per million (CPM) by dividing the RPK of a gene family by the total RPK in a sample. From CPM, percent abundance was obtained.

### Supplementary Tables

| Supplementary Table 1. Top 25 Gene Sets Enriched in CRC Samples from PRJNA413956 Dataset as Measured by Normalized Enrichment Scores. | | | |
|---|---|---|---|
| **GO ID** | **GO Term** | **\|NES\|** | **p.adjust** |
| GO:0042254 | GO_RIBOSOME_BIOGENESIS | 2.54 | 0.0021 |
| GO:0000779 | GO_CONDENSED_CHROMOSOME_CENTROMERIC_REGION | 2.50 | 0.0021 |
| GO:0034660 | GO_NCRNA_METABOLIC_PROCESS | 2.49 | 0.0021 |
| GO:0000793 | GO_CONDENSED_CHROMOSOME | 2.47 | 0.0021 |
| GO:0006260 | GO_DNA_REPLICATION | 2.47 | 0.0021 |
| GO:0098687 | GO_CHROMOSOMAL_REGION | 2.47 | 0.0021 |
| GO:0140014 | GO_MITOTIC_NUCLEAR_DIVISION | 2.46 | 0.0021 |
| GO:0022613 | GO_RIBONUCLEOPROTEIN_COMPLEX_BIOGENESIS | 2.46 | 0.0021 |
| GO:0034470 | GO_NCRNA_PROCESSING | 2.46 | 0.0021 |
| GO:0016072 | GO_RRNA_METABOLIC_PROCESS | 2.45 | 0.0021 |
| GO:0006261 | GO_DNA_DEPENDENT_DNA_REPLICATION | 2.45 | 0.0021 |
| GO:0007059 | GO_CHROMOSOME_SEGREGATION | 2.45 | 0.0021 |
| GO:0000819 | GO_SISTER_CHROMATID_SEGREGATION | 2.45 | 0.0021 |
| GO:0000775 | GO_CHROMOSOME_CENTROMERIC_REGION | 2.44 | 0.0021 |
| GO:0000070 | GO_MITOTIC_SISTER_CHROMATID_SEGREGATION | 2.44 | 0.0021 |
| GO:0071103 | GO_DNA_CONFORMATION_CHANGE | 2.43 | 0.0021 |
| GO:0098813 | GO_NUCLEAR_CHROMOSOME_SEGREGATION | 2.39 | 0.0021 |
| GO:0000776 | GO_KINETOCHORE | 2.38 | 0.0021 |
| GO:0030684 | GO_PRERIBOSOME | 2.37 | 0.0021 |
| GO:0051983 | GO_REGULATION_OF_CHROMOSOME_SEGREGATION | 2.37 | 0.0021 |
| GO:0048285 | GO_ORGANELLE_FISSION | 2.37 | 0.0021 |
| GO:0006405 | GO_RNA_EXPORT_FROM_NUCLEUS | 2.36 | 0.0021 |
| GO:0000075 | GO_CELL_CYCLE_CHECKPOINT | 2.36 | 0.0021 |
| GO:0051783 | GO_REGULATION_OF_NUCLEAR_DIVISION | 2.35 | 0.0021 |
| GO:0034728 | GO_NUCLEOSOME_ORGANIZATION | 2.34 | 0.0021 |
| GO:0032200 | GO_TELOMERE_ORGANIZATION | 2.33 | 0.0021 |

| Supplementary Table 2. Top 25 Gene Sets Enriched in CMS1 Dataset as Measured by Normalized Enrichment Scores against CMS2. | | | |
|---|---|---|---|
| **GO ID** | **GO Term** | **\|NES\|** | **p.adjust** |
| GO:0034341 | GO_RESPONSE_TO_INTERFERON_GAMMA | 2.62 | 0.0022 |
| GO:0060333 | GO_INTERFERON_GAMMA_MEDIATED_SIGNALING_PATHWAY | 2.53 | 0.0022 |
| GO:0002250 | GO_ADAPTIVE_IMMUNE_RESPONSE | 2.49 | 0.0022 |
| GO:0007159 | GO_LEUKOCYTE_CELL_CELL_ADHESION | 2.40 | 0.0022 |
| GO:0051607 | GO_DEFENSE_RESPONSE_TO_VIRUS | 2.39 | 0.0022 |
| GO:0042110 | GO_T_CELL_ACTIVATION | 2.38 | 0.0022 |
| GO:0045088 | GO_REGULATION_OF_INNATE_IMMUNE_RESPONSE | 2.37 | 0.0022 |
| GO:0001909 | GO_LEUKOCYTE_MEDIATED_CYTOTOXICITY | 2.37 | 0.0022 |
| GO:0001906 | GO_CELL_KILLING | 2.36 | 0.0022 |
| GO:0034340 | GO_RESPONSE_TO_TYPE_I_INTERFERON | 2.36 | 0.0022 |
| GO:0031341 | GO_REGULATION_OF_CELL_KILLING | 2.36 | 0.0022 |
| GO:0050863 | GO_REGULATION_OF_T_CELL_ACTIVATION | 2.36 | 0.0022 |
| GO:0002449 | GO_LYMPHOCYTE_MEDIATED_IMMUNITY | 2.35 | 0.0022 |
| GO:0002228 | GO_NATURAL_KILLER_CELL_MEDIATED_IMMUNITY | 2.34 | 0.0022 |
| GO:1903039 | GO_POSITIVE_REGULATION_OF_LEUKOCYTE_CELL_CELL_ADHESION | 2.33 | 0.0022 |
| GO:0002703 | GO_REGULATION_OF_LEUKOCYTE_MEDIATED_IMMUNITY | 2.32 | 0.0022 |
| GO:0042098 | GO_T_CELL_PROLIFERATION | 2.32 | 0.0022 |
| GO:0002697 | GO_REGULATION_OF_IMMUNE_EFFECTOR_PROCESS | 2.32 | 0.0022 |
| GO:0032609 | GO_INTERFERON_GAMMA_PRODUCTION | 2.32 | 0.0022 |
| GO:0009615 | GO_RESPONSE_TO_VIRUS | 2.31 | 0.0022 |
| GO:0001818 | GO_NEGATIVE_REGULATION_OF_CYTOKINE_PRODUCTION | 2.30 | 0.0022 |
| GO:0098542 | GO_DEFENSE_RESPONSE_TO_OTHER_ORGANISM | 2.30 | 0.0022 |
| GO:0002237 | GO_RESPONSE_TO_MOLECULE_OF_BACTERIAL_ORIGIN | 2.30 | 0.0022 |
| GO:0050852 | GO_T_CELL_RECEPTOR_SIGNALING_PATHWAY | 2.30 | 0.0022 |
| GO:0001819 | GO_POSITIVE_REGULATION_OF_CYTOKINE_PRODUCTION | 2.30 | 0.0022 |

| **Supplementary Table 3. Top 25 Gene Sets Enriched in CMS1 Dataset as Measured by Normalized Enrichment Scores against CMS3.** | | | |
|---|---|---|---|
| **GO ID** | **GO Term** | **\|NES\|** | **p.adjust** |
| GO:0051607 | GO_DEFENSE_RESPONSE_TO_VIRUS | 2.38 | 0.0024 |
| GO:0034341 | GO_RESPONSE_TO_INTERFERON_GAMMA | 2.34 | 0.0024 |
| GO:0009615 | GO_RESPONSE_TO_VIRUS | 2.32 | 0.0024 |
| GO:0098542 | GO_DEFENSE_RESPONSE_TO_OTHER_ORGANISM | 2.28 | 0.0024 |
| GO:0071216 | GO_CELLULAR_RESPONSE_TO_BIOTIC_STIMULUS | 2.28 | 0.0024 |
| GO:0045088 | GO_REGULATION_OF_INNATE_IMMUNE_RESPONSE | 2.26 | 0.0024 |
| GO:0048002 | GO_ANTIGEN_PROCESSING_AND_PRESENTATION_OF_PEPTIDE_ANTIGEN | 2.25 | 0.0024 |
| GO:0034340 | GO_RESPONSE_TO_TYPE_I_INTERFERON | 2.25 | 0.0024 |
| GO:0030199 | GO_COLLAGEN_FIBRIL_ORGANIZATION | 2.24 | 0.0024 |
| GO:0032611 | GO_INTERLEUKIN_1_BETA_PRODUCTION | 2.23 | 0.0024 |
| GO:0019882 | GO_ANTIGEN_PROCESSING_AND_PRESENTATION | 2.22 | 0.0024 |
| GO:0042590 | GO_ANTIGEN_PROCESSING_AND_PRESENTATION_OF_EXOGENOUS_PEPTIDE_ ANTIGEN_VIA_MHC_CLASS_I | 2.22 | 0.0024 |
| GO:0032612 | GO_INTERLEUKIN_1_PRODUCTION | 2.21 | 0.0024 |
| GO:0060333 | GO_INTERFERON_GAMMA_MEDIATED_SIGNALING_PATHWAY | 2.20 | 0.0024 |
| GO:0002474 | GO_ANTIGEN_PROCESSING_AND_PRESENTATION_OF_PEPTIDE_ANTIGEN_ VIA_MHC_CLASS_I | 2.18 | 0.0024 |
| GO:0005201 | GO_EXTRACELLULAR_MATRIX_STRUCTURAL_CONSTITUENT | 2.18 | 0.0024 |
| GO:0043062 | GO_EXTRACELLULAR_STRUCTURE_ORGANIZATION | 2.18 | 0.0024 |
| GO:0001819 | GO_POSITIVE_REGULATION_OF_CYTOKINE_PRODUCTION | 2.18 | 0.0024 |
| GO:0050663 | GO_CYTOKINE_SECRETION | 2.18 | 0.0024 |
| GO:0005581 | GO_COLLAGEN_TRIMER | 2.17 | 0.0024 |
| GO:0032635 | GO_INTERLEUKIN_6_PRODUCTION | 2.17 | 0.0024 |
| GO:0070555 | GO_RESPONSE_TO_INTERLEUKIN_1 | 2.17 | 0.0024 |
| GO:0071706 | GO_TUMOR_NECROSIS_FACTOR_SUPERFAMILY_CYTOKINE_PRODUCTION | 2.15 | 0.0024 |
| GO:0071887 | GO_LEUKOCYTE_APOPTOTIC_PROCESS | 2.15 | 0.0024 |
| GO:0031349 | GO_POSITIVE_REGULATION_OF_DEFENSE_RESPONSE | 2.12 | 0.0024 |

| **Supplementary Table 4. Top 25 Gene Sets Enriched in CMS2 Dataset as Measured by Normalized Enrichment Scores against CMS1.** | | | |
|---|---|---|---|
| **GO ID** | **GO Term** | **\|NES\|** | **p.adjust** |
| GO:0042273 | GO_RIBOSOMAL_LARGE_SUBUNIT_BIOGENESIS | 2.10 | 0.0042 |
| GO:0140053 | GO_MITOCHONDRIAL_GENE_EXPRESSION | 2.04 | 0.0056 |
| GO:0032543 | GO_MITOCHONDRIAL_TRANSLATION | 1.97 | 0.0052 |
| GO:0042788 | GO_POLYSOMAL_RIBOSOME | 1.93 | 0.0035 |
| GO:0008033 | GO_TRNA_PROCESSING | 1.93 | 0.0052 |
| GO:0044391 | GO_RIBOSOMAL_SUBUNIT | 1.92 | 0.006 |
| GO:0042254 | GO_RIBOSOME_BIOGENESIS | 1.92 | 0.0079 |
| GO:0000462 | GO_MATURATION_OF_SSU_RRNA_FROM_TRICISTRONIC_RRNA_TRANSCRIPT_ SSU_RRNA_5_8S_RRNA_LSU_RRNA | 1.92 | 0.0037 |
| GO:0030490 | GO_MATURATION_OF_SSU_RRNA | 1.90 | 0.0063 |
| GO:0034660 | GO_NCRNA_METABOLIC_PROCESS | 1.90 | 0.0116 |
| GO:0006400 | GO_TRNA_MODIFICATION | 1.90 | 0.0044 |
| GO:0016072 | GO_RRNA_METABOLIC_PROCESS | 1.90 | 0.0063 |
| GO:0006399 | GO_TRNA_METABOLIC_PROCESS | 1.89 | 0.0059 |
| GO:0000184 | GO_NUCLEAR_TRANSCRIBED_MRNA_CATABOLIC_PROCESS_NONSENSE_ MEDIATED_DECAY | 1.88 | 0.0051 |
| GO:0006270 | GO_DNA_REPLICATION_INITIATION | 1.88 | 0.0060 |
| GO:0003735 | GO_STRUCTURAL_CONSTITUENT_OF_RIBOSOME | 1.88 | 0.0056 |
| GO:0030684 | GO_PRERIBOSOME | 1.88 | 0.0042 |
| GO:0042274 | GO_RIBOSOMAL_SMALL_SUBUNIT_BIOGENESIS | 1.88 | 0.004 |
| GO:0070129 | GO_REGULATION_OF_MITOCHONDRIAL_TRANSLATION | 1.87 | 0.0097 |
| GO:0034470 | GO_NCRNA_PROCESSING | 1.85 | 0.0097 |
| GO:0005736 | GO_RNA_POLYMERASE_I_COMPLEX | 1.84 | 0.0087 |
| GO:0001510 | GO_RNA_METHYLATION | 1.84 | 0.0043 |
| GO:0098798 | GO_MITOCHONDRIAL_PROTEIN_COMPLEX | 1.84 | 0.0073 |
| GO:0015934 | GO_LARGE_RIBOSOMAL_SUBUNIT | 1.82 | 0.005 |
| GO:0022626 | GO_CYTOSOLIC_RIBOSOME | 1.81 | 0.005 |

| Supplementary Table 5.Top 25 Gene Sets Enriched in CMS2 Dataset as Measured by Normalized Enrichment Scores against CMS3. | | | |
|---|---|---|---|
| **GO ID** | **GO Term** | **\|NES\|** | **p.adjust** |
| GO:0034660 | GO_NCRNA_METABOLIC_PROCESS | 2.48 | 0.0029 |
| GO:0042254 | GO_RIBOSOME_BIOGENESIS | 2.46 | 0.0029 |
| GO:0034470 | GO_NCRNA_PROCESSING | 2.45 | 0.0029 |
| GO:0022613 | GO_RIBONUCLEOPROTEIN_COMPLEX_BIOGENESIS | 2.44 | 0.0029 |
| GO:0016072 | GO_RRNA_METABOLIC_PROCESS | 2.43 | 0.0029 |
| GO:0030684 | GO_PRERIBOSOME | 2.41 | 0.0029 |
| GO:0006270 | GO_DNA_REPLICATION_INITIATION | 2.40 | 0.0029 |
| GO:0032543 | GO_MITOCHONDRIAL_TRANSLATION | 2.31 | 0.0029 |
| GO:0006399 | GO_TRNA_METABOLIC_PROCESS | 2.29 | 0.0029 |
| GO:0032040 | GO_SMALL_SUBUNIT_PROCESSOME | 2.28 | 0.0029 |
| GO:0140053 | GO_MITOCHONDRIAL_GENE_EXPRESSION | 2.28 | 0.0029 |
| GO:0042273 | GO_RIBOSOMAL_LARGE_SUBUNIT_BIOGENESIS | 2.21 | 0.0029 |
| GO:0008033 | GO_TRNA_PROCESSING | 2.20 | 0.0029 |
| GO:0006415 | GO_TRANSLATIONAL_TERMINATION | 2.16 | 0.0029 |
| GO:0007143 | GO_FEMALE_MEIOTIC_NUCLEAR_DIVISION | 2.14 | 0.0029 |
| GO:0120114 | GO_SM_LIKE_PROTEIN_FAMILY_COMPLEX | 2.11 | 0.0029 |
| GO:0000313 | GO_ORGANELLAR_RIBOSOME | 2.09 | 0.0029 |
| GO:0044391 | GO_RIBOSOMAL_SUBUNIT | 2.08 | 0.0029 |
| GO:0003688 | GO_DNA_REPLICATION_ORIGIN_BINDING | 2.08 | 0.0029 |
| GO:0070126 | GO_MITOCHONDRIAL_TRANSLATIONAL_TERMINATION | 2.08 | 0.0029 |
| GO:0140101 | GO_CATALYTIC_ACTIVITY_ACTING_ON_A_TRNA | 2.08 | 0.0029 |
| GO:0071826 | GO_RIBONUCLEOPROTEIN_COMPLEX_SUBUNIT_ORGANIZATION | 2.02 | 0.0029 |
| GO:0000375 | GO_RNA_SPLICING_VIA_TRANSESTERIFICATION_REACTIONS | 2.02 | 0.0029 |
| GO:0015934 | GO_LARGE_RIBOSOMAL_SUBUNIT | 2.02 | 0.0029 |
| GO:0000184 | GO_NUCLEAR_TRANSCRIBED_MRNA_CATABOLIC_PROCESS_NONSENSE_ MEDIATED_DECAY | 2.02 | 0.0029 |

| **Supplementary Table 6. Top 25 Gene Sets Enriched in CMS3 Dataset as Measured by Normalized Enrichment Scores against CMS1.** | | | |
|---|---|---|---|
| **GO ID** | **GO Term** | **\|NES\|** | **p.adjust** |
| GO:0009812 | GO_FLAVONOID_METABOLIC_PROCESS | 2.22 | 0.0032 |
| GO:0072329 | GO_MONOCARBOXYLIC_ACID_CATABOLIC_PROCESS | 2.22 | 0.0037 |
| GO:0009062 | GO_FATTY_ACID_CATABOLIC_PROCESS | 2.19 | 0.0037 |
| GO:0006805 | GO_XENOBIOTIC_METABOLIC_PROCESS | 2.17 | 0.0037 |
| GO:0034440 | GO_LIPID_OXIDATION | 2.16 | 0.0037 |
| GO:0006635 | GO_FATTY_ACID_BETA_OXIDATION | 2.14 | 0.0035 |
| GO:0003707 | GO_STEROID_HORMONE_RECEPTOR_ACTIVITY | 2.11 | 0.0034 |
| GO:0006631 | GO_FATTY_ACID_METABOLIC_PROCESS | 2.07 | 0.0047 |
| GO:0044242 | GO_CELLULAR_LIPID_CATABOLIC_PROCESS | 2.06 | 0.0043 |
| GO:0006063 | GO_URONIC_ACID_METABOLIC_PROCESS | 2.04 | 0.0032 |
| GO:0034308 | GO_PRIMARY_ALCOHOL_METABOLIC_PROCESS | 2.04 | 0.0035 |
| GO:0016408 | GO_C_ACYLTRANSFERASE_ACTIVITY | 1.99 | 0.0032 |
| GO:0005903 | GO_BRUSH_BORDER | 1.99 | 0.0037 |
| GO:0045277 | GO_RESPIRATORY_CHAIN_COMPLEX_IV | 1.99 | 0.0032 |
| GO:0001972 | GO_RETINOIC_ACID_BINDING | 1.93 | 0.0064 |
| GO:0016614 | GO_OXIDOREDUCTASE_ACTIVITY_ACTING_ON_CH_OH_GROUP_OF_DONORS | 1.92 | 0.0037 |
| GO:0034754 | GO_CELLULAR_HORMONE_METABOLIC_PROCESS | 1.92 | 0.0036 |
| GO:0032787 | GO_MONOCARBOXYLIC_ACID_METABOLIC_PROCESS | 1.91 | 0.0060 |
| GO:0071280 | GO_CELLULAR_RESPONSE_TO_COPPER_ION | 1.91 | 0.0081 |
| GO:0004879 | GO_NUCLEAR_RECEPTOR_ACTIVITY | 1.90 | 0.0052 |
| GO:0016042 | GO_LIPID_CATABOLIC_PROCESS | 1.90 | 0.0050 |
| GO:0033559 | GO_UNSATURATED_FATTY_ACID_METABOLIC_PROCESS | 1.90 | 0.0035 |
| GO:0015701 | GO_BICARBONATE_TRANSPORT | 1.89 | 0.0068 |
| GO:0033293 | GO_MONOCARBOXYLIC_ACID_BINDING | 1.89 | 0.0055 |
| GO:0033540 | GO_FATTY_ACID_BETA_OXIDATION_USING_ACYL_COA_OXIDASE | 1.88 | 0.0093 |

| **Supplementary Table 7. Top 25 Gene Sets Enriched in CMS3 Dataset as Measured by Normalized Enrichment Scores against CMS2.** | | | |
|---|---|---|---|
| **GO ID** | **GO Term** | **\|NES\|** | **p.adjust** |
| GO:0006063 | GO_URONIC_ACID_METABOLIC_PROCESS | 2.09 | 0.0029 |
| GO:0071294 | GO_CELLULAR_RESPONSE_TO_ZINC_ION | 2.07 | 0.0029 |
| GO:0015020 | GO_GLUCURONOSYLTRANSFERASE_ACTIVITY | 2.06 | 0.0029 |
| GO:0009812 | GO_FLAVONOID_METABOLIC_PROCESS | 2.05 | 0.0029 |
| GO:0071276 | GO_CELLULAR_RESPONSE_TO_CADMIUM_ION | 1.98 | 0.0029 |
| GO:0010043 | GO_RESPONSE_TO_ZINC_ION | 1.96 | 0.0029 |
| GO:0006805 | GO_XENOBIOTIC_METABOLIC_PROCESS | 1.96 | 0.0029 |
| GO:0034754 | GO_CELLULAR_HORMONE_METABOLIC_PROCESS | 1.96 | 0.0029 |
| GO:0004745 | GO_RETINOL_DEHYDROGENASE_ACTIVITY | 1.94 | 0.0029 |
| GO:0005496 | GO_STEROID_BINDING | 1.92 | 0.0029 |
| GO:0030258 | GO_LIPID_MODIFICATION | 1.91 | 0.0029 |
| GO:0016042 | GO_LIPID_CATABOLIC_PROCESS | 1.91 | 0.0029 |
| GO:0006635 | GO_FATTY_ACID_BETA_OXIDATION | 1.91 | 0.0029 |
| GO:0034440 | GO_LIPID_OXIDATION | 1.89 | 0.0029 |
| GO:0006654 | GO_PHOSPHATIDIC_ACID_BIOSYNTHETIC_PROCESS | 1.87 | 0.0029 |
| GO:0007586 | GO_DIGESTION | 1.86 | 0.0029 |
| GO:0006066 | GO_ALCOHOL_METABOLIC_PROCESS | 1.85 | 0.0029 |
| GO:0042572 | GO_RETINOL_METABOLIC_PROCESS | 1.83 | 0.0047 |
| GO:0001972 | GO_RETINOIC_ACID_BINDING | 1.82 | 0.0047 |
| GO:0016408 | GO_C_ACYLTRANSFERASE_ACTIVITY | 1.81 | 0.0078 |
| GO:0032411 | GO_POSITIVE_REGULATION_OF_TRANSPORTER_ACTIVITY | 1.81 | 0.0029 |
| GO:0005319 | GO_LIPID_TRANSPORTER_ACTIVITY | 1.81 | 0.0029 |
| GO:0044242 | GO_CELLULAR_LIPID_CATABOLIC_PROCESS | 1.80 | 0.0029 |
| GO:0008028 | GO_MONOCARBOXYLIC_ACID_TRANSMEMBRANE_TRANSPORTER_ACTIVITY | 1.79 | 0.0062 |
| GO:0042445 | GO_HORMONE_METABOLIC_PROCESS | 1.79 | 0.0029 |
